# Anthocyanin-associated cellular programs underlying terroir variation in Cabernet Sauvignon grape berry revealed by SEED-based deconvolution

**DOI:** 10.64898/2026.06.05.730035

**Authors:** Xinhao Hu, Yu Tang, Fulan Deng, Zhiyu Chen, Guwei Tang, Xu Yan, Zengqiang Xia, Henry H. Y. Tong, Jicheng Zhan, Xin Zou, Jie Hao

**Affiliations:** College of Food Science and Nutritional Engineering, Beijing Key Laboratory of Viticulture and Enology, China Agricultural University, Tsinghua East Road 17, HaidianDistrict, Beijing 100083, China; Shanghai Key Laboratory of Plant Functional Genomics and Resources, Shanghai Chenshan Botanical Garden, and Chenshan Science Research Center, CAS Center for Excellence in Molecular Plant Sciences (CEMPS), Chinese Academy of Sciences (CAS), Shanghai, China; Centre of AI-driven Drug Discovery, Faculty of Applied Sciences, Macao Polytechnic University, Macau, Macau SAR; Institute of Clinical Science, Zhongshan Hospital, Fudan University, Shanghai, China; Jishou University, Jishou, Hunan, China; National Center for Translational Medicine (Shanghai) Linyi Branch & Institute of Translational Medicine Linyi University; School of Medicine, Linyi University, Shuangling Road, Linyi, Shandong, China; Digital Diagnosis and Treatment Innovation Center for Cancer, Institute of Translational Medicine, Shanghai Jiao Tong University, Shanghai, China

**Keywords:** scRNA-seq, bulk RNA-seq, deconvolution

## Abstract

Plant tissues consist of diverse cell populations that collectively contribute to development, metabolism, environmental responses, and phenotype formation. Although single-cell and single-nucleus RNA sequencing have greatly advanced the study of plant cellular heterogeneity, their application to large sample cohorts remains limited by cost, technical complexity, tissue dissociation constraints, and throughput. In contrast, bulk RNA-seq datasets have accumulated extensively across plant species, tissues, developmental stages, and environmental conditions, yet the celltype-level information embedded in these datasets remains difficult to resolve because plant-oriented deconvolution frameworks are still lacking. Existing deconvolution methods have largely been developed in mammalian systems and have not been systematically optimized for plant transcriptomic features, leaving their applicability under plant-specific constraints unclear. Here, we present SEED, an adaptive deconvolution framework optimized for plant transcriptomic data. SEED integrates candidate reference-template construction with seven deconvolution strategies and automatically identifies an optimal combination for a given dataset. In grapevine simulated benchmarking, SEED showed its clearest advantage under low-replication conditions and remained broadly competitive, rather than uniformly dominant, when larger pseudo-bulk sample sizes were evaluated. SEED further performed robustly in public *Arabidopsis thaliana* and *Nicotiana tabacum* datasets. Finally, we applied SEED to bulk RNA-seq data generated in this study from *Vitis vinifera cv*. Cabernet Sauvignon berries collected from Yinchuan and Yantai, identifying terroir-associated cell subtypes and coordinated celltype interaction patterns. Together, these results establish SEED as a practical framework for plant transcriptome deconvolution and provide a new tool for dissecting cellular heterogeneity associated with environmental adaptation and phenotype formation in plants.

## Introduction

Plant organs are composed of diverse specialized cell populations whose relative abundance and transcriptional states collectively shape tissue development, physiological function, and phenotype formation^1,2^. The organization and coordination of these cell populations underlie key biological processes, including growth, differentiation, metabolic regulation, and environmental responses^3–5^. A celltype-resolved understanding of plant tissues is therefore essential for elucidating how developmental programs and external cues are integrated to generate phenotypic variation^6^.

Single-cell and single-nucleus transcriptomic approaches have greatly expanded the capacity to study cellular heterogeneity in plants^7,8^, and their use is increasing across roots, leaves, reproductive tissues, and fruits^9–11^. At the same time, plant single-cell analysis remains constrained by technical complexity^12^, tissue-specific dissociation challenges^13^, cost and throughput^14^, making it difficult to apply routinely to large experimental cohorts. In contrast, bulk RNA-seq remains far more abundant across species, tissues, and environmental conditions, providing broader contextual coverage and greater practical scalability than single-cell approaches^15^. Accordingly, reference-based deconvolution offers an attractive strategy for combining the cellular resolution of single-cell data with the scale and practicality of bulk transcriptomics^16,17^.

Despite this opportunity, most deconvolution frameworks have been developed and benchmarked in biomedical contexts rather than in plant systems^18,19^. Their assumptions, reference strategies, and optimization procedures have not been systematically tailored to plant transcriptomic data, and their performance in plants remains unclear^20,21^. As a result, plant studies still lack a robust and broadly validated deconvolution framework capable of accurately estimating celltype composition from bulk transcriptomes.

A major limitation of plant transcriptome deconvolution lies in the reference template itself. Accurate deconvolution depends on how well the reference represents the target bulk sample^21,22^. In plants, this remains particularly challenging because single-cell references are still sparse across species, tissues, developmental stages, and environmental conditions^6,23,24^, limiting the construction of representative templates. The problem is further compounded by the way plant single-cell data are generated: most workflows rely on enzymatic digestion of the cell wall to produce protoplasts, or alternatively on nucleus extraction, and neither strategy recovers all cell types equally well^25,26^. Moreover, protoplasting can perturb native transcriptional states, introducing bias into the resulting reference profiles^27^. These limitations become especially consequential in plant studies, which are often conducted with small sample sizes and under strong environmental variation. Under such conditions, deconvolution performance becomes highly sensitive to template construction^22^, annotation granularity^28^ and parameter choice^29^.Together, these features indicate that plant bulk RNA-seq deconvolution requires dedicated optimization of reference design, sample matching and robustness under small-sample conditions, rather than a direct transfer of frameworks developed for animal systems.

In grapevine research, terroir is a central concept for understanding how environmental variation shapes fruit composition and quality^30^. Even within the same cultivar, differences in climate, soil and cultivation context can lead to substantial variation in berry composition and wine style^31,32^. However, whether terroir-associated variation is reflected at the level of tissue cellular composition remains largely unknown. Here we present SEED, an adaptive deconvolution framework developed for plant transcriptomic data. Building on our previous DECEPTICONx^33^ framework for biomedical systems, we optimized the workflow for plant data, benchmarked it in grapevine simulated datasets, validated it in public *Arabidopsis thaliana*^34^ and *Nicotiana tabacum*^35^ datasets, and applied it to grapevine bulk RNA-seq data generated in this study from Yantai and Yinchuan production regions. Collectively, our results establish SEED as a robust framework for plant transcriptome deconvolution and provide a basis for resolving phenotype-associated cellular heterogeneity in plants.

## Results

### 1. SEED framework and benchmarking in grapevine simulated datasets

Inspired by our previously developed DECEPTICONx framework for biomedical systems, we adapted and optimized the workflow for plant transcriptomic data, resulting in SEED. SEED is a context-optimized deconvolution framework that takes single-cell or single-nucleus RNA-seq data and bulk RNA-seq data as input. In this study, we use the term Reference Template to refer to celltype-resolved reference expression profiles, typically represented as a matrix of molecular signatures for distinct cell types, which provide the prior information required for bulk transcriptome deconvolution. Within SEED, candidate Reference Templates are generated using complementary strategies, including Monocle3^36–38^-based aggregation and BayesPrism^39^-derived reference profiles. These templates are then evaluated in combination with seven deconvolution methods—CIBERSORT^40^, quanTIseq^41^, EPIC^42^, DeconRNASeq^43^, MuSiC^18^, SCDC^44^, and MCP-counter^45^—allowing SEED to automatically identify the optimal pairing of template construction and deconvolution strategy for a given dataset(Fig.1a).

**Figure 1.**
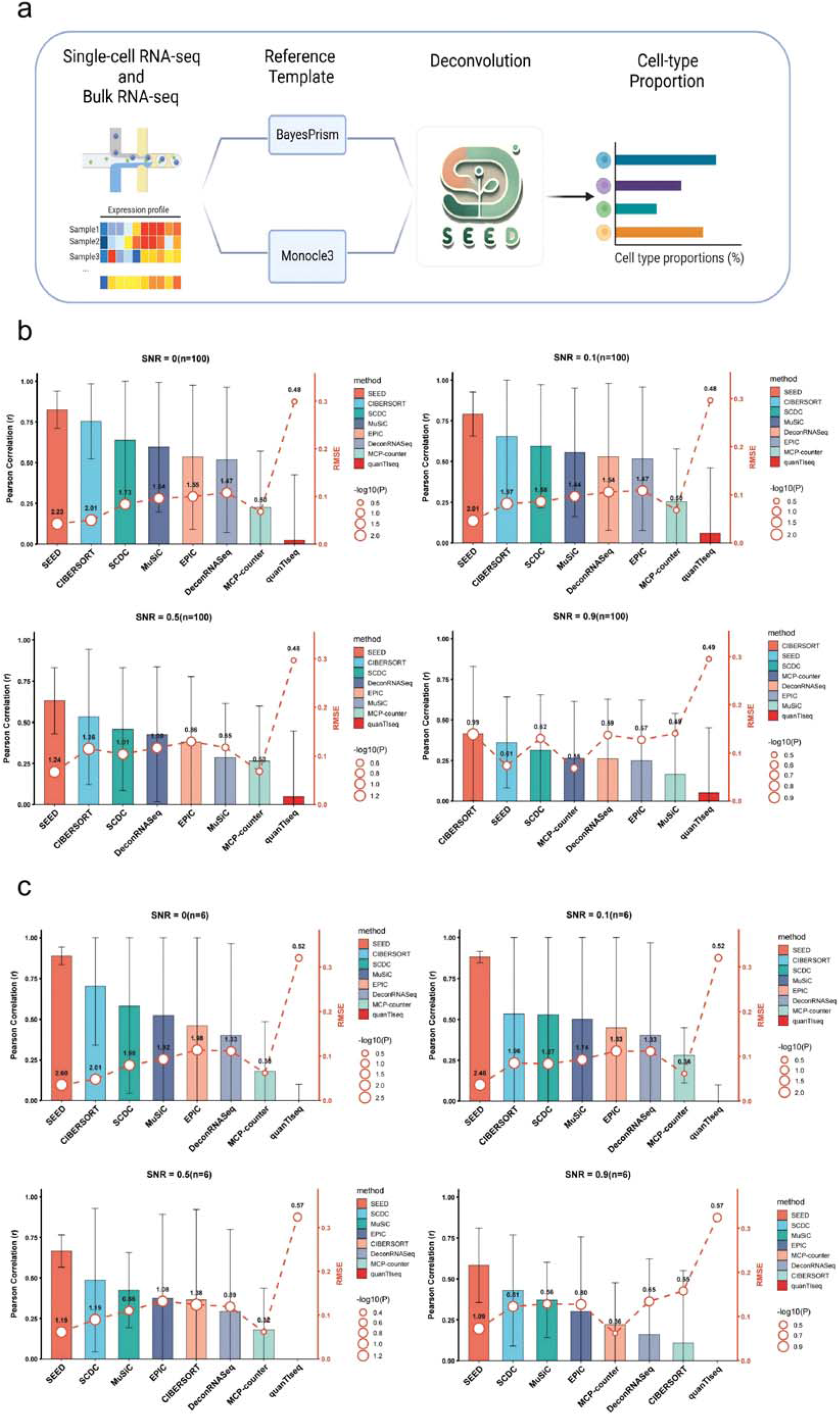
Overview of the SEED workflow and simulated grape pseudo-bulk benchmarking. (a) SEED integrates single-cell/single-nucleus and bulk RNA-seq inputs, constructs candidate reference templates using BayesPrism and Monocle3, and estimates celltype proportions through adaptive deconvolution. (b) Benchmarking under the larger n = 100 condition across SNR = 0, 0.1, 0.5, and 0.9. Bars show Pearson correlation (r; higher values indicate better agreement with the expected proportions), red lines indicate RMSE (lower values indicate smaller estimation error), and red circles/labels indicate -log10(P) (larger values indicate stronger statistical support). (c) Benchmarking under the low-replication n = 6 condition across SNR = 0, 0.1, 0.5, and 0.9. Under this more stringent setting, SEED ranked first by Pearson correlation across all four SNR levels and showed the strongest or near-strongest overall performance when RMSE and -log10(P) were considered together.

To evaluate SEED in a plant-relevant setting, we generated grape pseudo-bulk RNA-seq datasets as simulated datasets from *Vitis vinifera cv*. Cabernet Sauvignon single-nucleus transcriptomic data generated in this study^46^. We first evaluated a larger pseudo-bulk setting with n = 100 samples to examine whether SEED maintained competitive performance under relatively well-sampled conditions (Fig. 1b). In this simulation, the SNR parameter was used to mimic increasing noise perturbation in realistic bulk RNA-seq data: SNR = 0 represents the no-added-noise pseudo-bulk condition, whereas SNR = 0.1, 0.5, and 0.9 represent progressively stronger deviations from the idealized mixture. Under n = 100, SEED achieved the best overall performance at SNR = 0 and 0.1, but its ranking fluctuated at higher noise levels. These results indicate that algorithmic superiority under abundant pseudo-bulk sampling is context-dependent. Because plant transcriptomic studies are frequently constrained by field sampling, material availability, and sequencing cost, we therefore further tested the more stringent low-replication condition with n = 6 samples under the same four SNR levels (Fig. 1c). Under this low-replication setting, SEED achieved the strongest overall performance, combining high Pearson correlation (r), low root mean square error (RMSE), and strong statistical support (-log10(P)) across the tested noise levels. These results indicate that SEED is particularly advantageous when biological replication is limited, while the n = 100 benchmark provides a complementary assessment showing that its performance remains broadly competitive but may fluctuate under less restrictive sampling conditions.

### 2. SEED exhibits robust performance across public Arabidopsis thaliana and Nicotiana tabacum datasets

Following the grapevine simulated benchmark, we next asked whether the strong performance of SEED could be extended to real plant datasets. To address this, we evaluated SEED in public *Arabidopsis thaliana* and *Nicotiana tabacum* datasets representing distinct species backgrounds and experimental contexts.

In the *Arabidopsis thaliana* dataset, SEED achieved the best performance across all three evaluation metrics, including correlation, RMSE, and statistical significance (Fig. 2a,b). This result indicates that SEED can maintain high deconvolution accuracy in an external plant dataset and supports its applicability beyond the grape benchmarking system.

**Figure 2.**
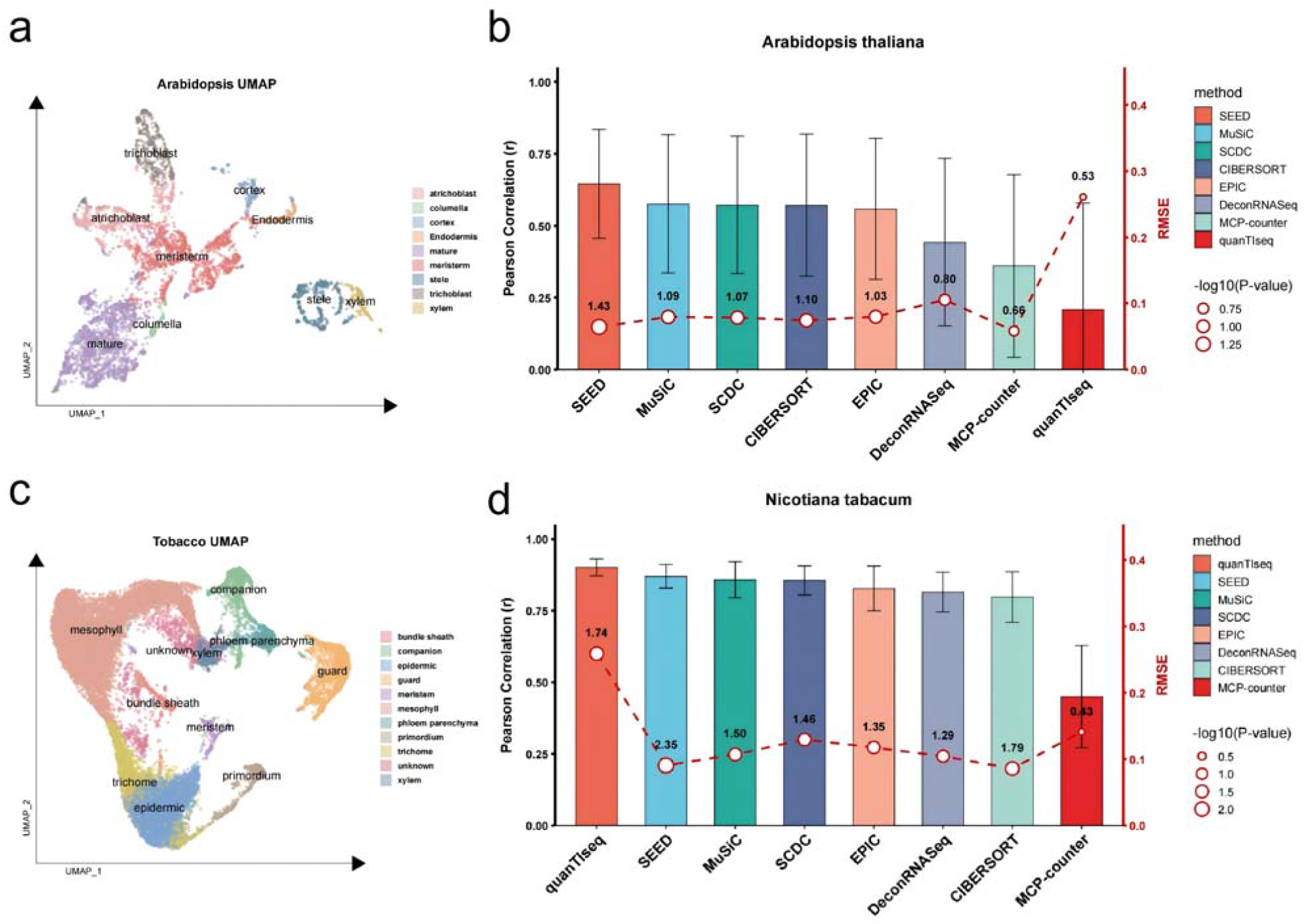
Cross-species evaluation of SEED in public Arabidopsis thaliana and Nicotiana tabacum datasets. (a) UMAP of annotated *Arabidopsis* scRNA-seq. (b) Deconvolution performance in Arabidopsis. SEED showed the strongest overall performance across the displayed metrics, ranking first by Pearson correlation and showing high statistical support while maintaining low estimation error. (c) UMAP of annotated Tobacco(*Nicotiana tabacum*) scRNA-seq. (d) Deconvolution performance in Nicotiana tabacum. quanTIseq ranked first by Pearson correlation, whereas SEED ranked second by correlation and showed strong statistical support, indicating stable performance across a distinct species background. In panels b and d, bars show Pearson correlation (r; higher is better), red lines indicate RMSE (lower is better), and red circles/labels indicate - log10(P) (higher is better).

In the *Nicotiana tabacum* dataset, SEED also showed strong overall performance (Fig. 2c,d). Although it ranked second in correlation, the difference from the top-performing method (quanTiseq) was small, while its performance across the other evaluation metrics remained competitive. This pattern suggests that, although the relative ranking of individual methods may vary across datasets, SEED retains stable and accurate performance in independent plant systems. Together, these results indicate that the utility of SEED is not restricted to the grapevine benchmark and that the framework can generalize to real plant datasets with distinct biological and technical backgrounds.

### 3. Single-nucleus profiling and celltype annotation of grapevine berry support downstream deconvolution

To provide a biologically relevant reference for downstream deconvolution, we performed single-nucleus RNA sequencing (snRNA-seq) on mature berry tissues of Vitis vinifera cv. Cabernet Sauvignon collected from Yantai (Fig. 3a). Following the experimental workflow illustrated in Fig. 3a, seeds were first removed from Cabernet Sauvignon berries to enrich for skin and flesh components. Fresh tissue was then processed for nuclei extraction under low-temperature conditions, and the purified nuclei suspension was loaded into the 10x Genomics microfluidic system to generate single-nucleus droplets. Barcoded cDNA libraries were constructed from captured nuclei and sequenced to obtain nucleus-resolved transcriptomes. After read processing, quality control, dimensionality reduction, and clustering, the resulting expression matrix was used to define transcriptionally distinct grapevine berry cell clusters. Eight clusters were resolved by UMAP and represented the major cellular components captured from berry tissues (Fig. 3b). Although this dataset was generated from a single biological sample and was not intended to constitute a comprehensive single-nucleus atlas, it provided a practical celltype reference framework for subsequent bulk deconvolution analysis.

**Figure 3.**
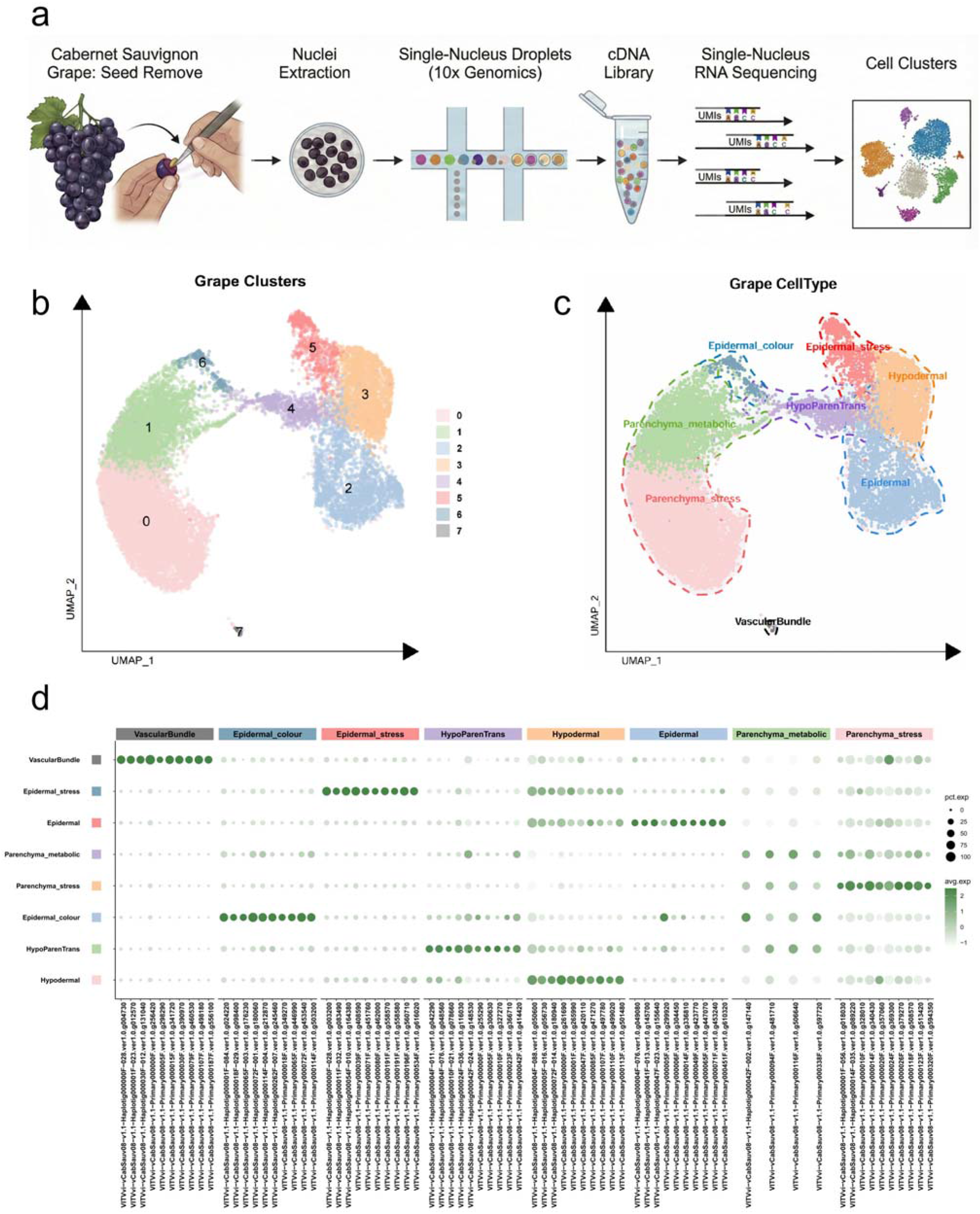
Single-nucleus RNA-seq workflow and celltype annotation of Cabernet Sauvignon grape berry. (a) Experimental workflow from seed removal, nuclei extraction, 10x Genomics droplet generation, cDNA library construction, and single-nucleus RNA sequencing to cell clustering. (b) UMAP visualization of transcriptionally resolved grapevine berry clusters before celltype annotation. (c) UMAP visualization of the eight annotated grapevine berry celltypes, including Epidermal, Epidermal_colour, Epidermal_stress, Hypodermal, HypoParenTrans, Parenchyma_stress, VascularBundle, and Parenchyma_metabolic. (d) Dotplot of the top 10 marker genes. Dot size represents the proportion of nuclei expressing each marker gene, and color intensity represents scaled average expression.

To assign biological identities to these clusters, we integrated cluster-specific marker genes with pathway-related functional information and previously reported marker evidence from grape and related plant species. Based on these combined criteria, the eight clusters were annotated as Epidermal, Epidermal_colour, Epidermal_stress, Hypodermal, HypoParenTrans, Parenchyma_stress, VascularBundle, and Parenchyma_metabolic (Fig. 3c,d). These annotations captured both structural and functional diversity within berry tissues, including outer tissue-associated populations, transport-related cells, and metabolic or stress-associated cell states.

Importantly, this reference was not used as an endpoint in itself, but as the basis for interpreting bulk transcriptomic variation at celltype resolution. By defining the principal cell populations represented in grape berry tissues, the snRNA-seq dataset established the cellular framework required for SEED-based deconvolution and enabled downstream interrogation of region-associated cellular variation.

### 4. SEED reveals terroir-associated cell subtypes and coordinated cellular interactions in grapevine berry tissues

We next applied SEED to bulk RNA-seq data from mature *Vitis vinifera cv*. Cabernet Sauvignon berries collected from two geographically distinct production regions, Yantai and Yinchuan, to determine whether terroir-associated variation could be resolved at the level of cellular composition. SEED deconvolution identified multiple cell populations showing significant regional differences, including Epidermal, Epidermal_colour, Epidermal_stress, Hypodermal, HypoParenTrans, Parenchyma_stress, and VascularBundle, whereas Parenchyma_metabolic did not differ significantly between the two regions (Fig. 4a). Quantitative deconvolution further showed that the two regions differed in overall cellular composition (Fig. 4b). Yantai samples were enriched in Epidermal, Hypodermal, HypoParenTrans, and Parenchyma_stress populations, whereas Yinchuan samples showed higher proportions of Epidermal_colour, Epidermal_stress, and VascularBundle populations. In contrast, Parenchyma_metabolic remained comparatively stable between regions, suggesting that terroir-associated effects are concentrated in specific cellular compartments rather than distributed uniformly across the tissue.

**Figure 4.**
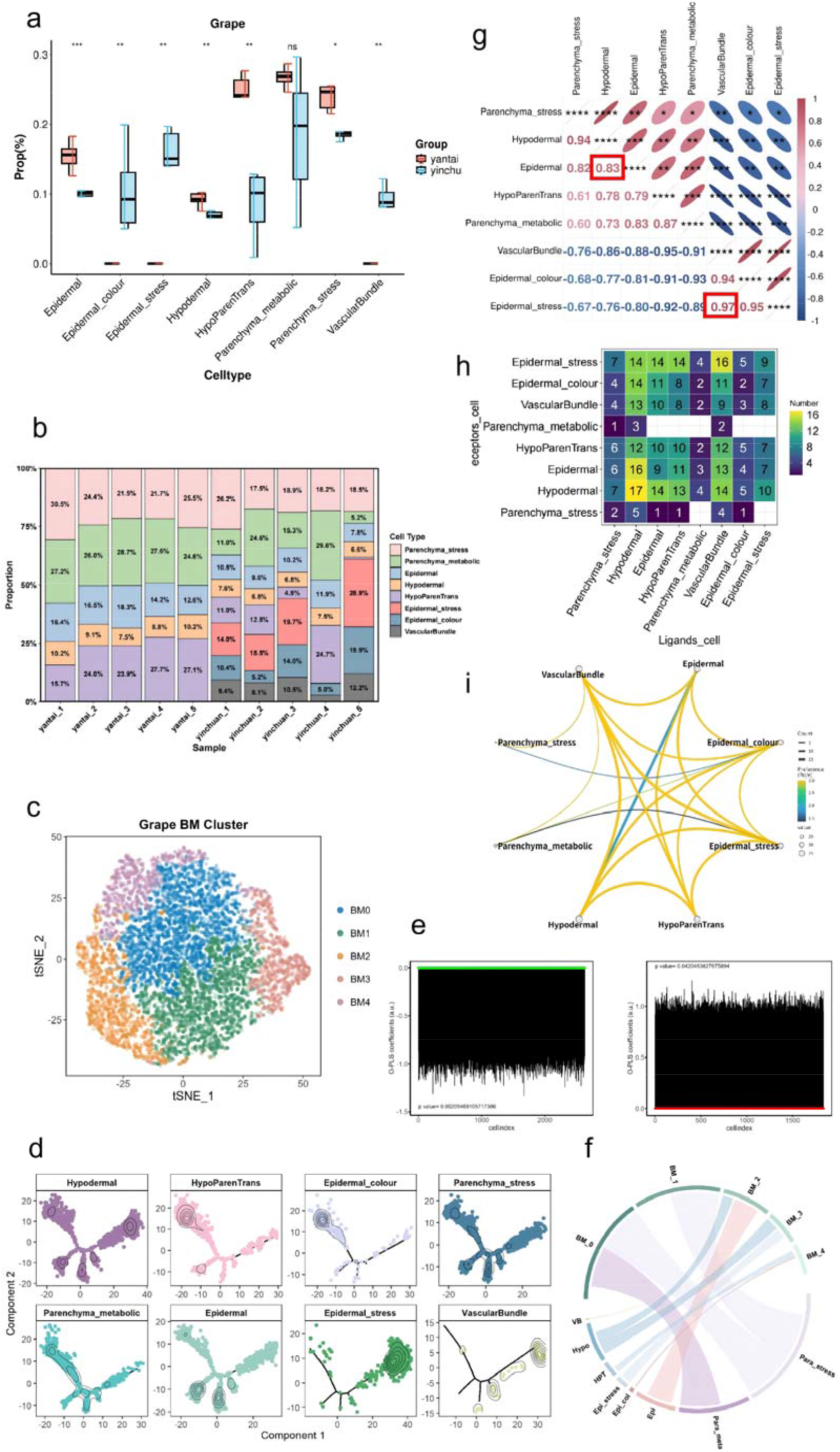
SEED resolves terroir-associated cellular variation and coordinated celltype relationships in Cabernet Sauvignon grape berries. (a) Regional differences in SEED-inferred celltype proportions between Yantai and Yinchuan. Significance levels are indicated above each cell type; ns denotes not significant. (b) Stacked bar plot showing the SEED-inferred cellular composition of each bulk RNA-seq sample, highlighting region-associated enrichment of Epidermal, Hypodermal, HypoParenTrans, and Parenchyma_stress in Yantai and enrichment of Epidermal_colour, Epidermal_stress, and VascularBundle in Yinchuan. (c) t-SNE visualization of BLOOM-reconstructed grape single-cell expression profiles. (d) OPLS-DA loading plots showing phenotype associations of BLOOM-derived grape celltypes. Top: Yantai; bottom: Yinchuan. (e) Pseudotime trajectory of grape cell expression profiles after BLOOM reconstruction, colored by original cell types. (f) Correspondence between original and BLOOM clusters in the grape dataset. Upper ring: BLOOM clusters; lower ring: original celltypes; chord width reflects cell numbers; only significant relationships (p < 0.05) are shown. (g) Correlation matrix of inferred celltype proportions. The highlighted pairs indicate strong positive correlations between Epidermal and Hypodermal (r = 0.83, p < 0.05) and between Epidermal_stress and VascularBundle (r = 0.97, p < 0.05). (h) Heatmap showing the number of ligand-receptor communication events between sender and receiver cell populations. (i) Cell-cell communication network based on the observed-to-expected interaction preference ratio (Ro/e). Only interactions with Ro/e > 1 are shown, indicating preferential communication beyond that expected from celltype abundance alone. Color represents the magnitude of the Ro/e ratio, with more yellow colors indicating higher Ro/e values. This analysis supports preferential interactions between Epidermal-Hypodermal and Epidermal_stress-VascularBundle cell pairs.

Additional analysis using BLOOM—our recently developed framework for Bulk and singLe-cell transcriptOmics integratiOn for Plant PhenoMics—further supported the regional patterns identified by SEED. In this grapevine dataset, BLOOM associated Epidermal with the Yantai phenotype and Epidermal_stress with the Yinchuan phenotype (Fig. 4c-f), consistent with the region-associated cellular shifts inferred by SEED. This concordance supports the interpretation that the regional signals captured by SEED reflect biologically meaningful variation rather than reference-specific bias alone.

To further examine whether these region-associated cell populations varied in a coordinated manner, we performed correlation analysis on the celltype proportion estimates generated by SEED. This analysis revealed a strong positive correlation between Epidermal and Hypodermal (r = 0.83, p < 0.05), as well as between Epidermal_stress and VascularBundle (r = 0.97, p < 0.05) (Fig. 4g). These results suggest that the major terroir-associated cell populations identified by SEED do not vary independently, but instead may respond in coordinated cellular modules.

We next performed cell-cell communication analysis on the snRNA-seq reference using PlantPhoneDB^47^. Consistent with the correlation patterns derived from the SEED-inferred celltype proportions, Epidermal and Hypodermal, as well as Epidermal_stress and VascularBundle, showed frequent communication events (Fig. 4h). To further exclude the possibility that these signals were driven simply by differences in cell abundance, we evaluated preferential communication using the observed-to-expected ratio (Ro/e)^48^, in which the observed number of ligand-receptor interactions is compared with the expected number based on celltype abundance. Both Epidermal-Hypodermal and Epidermal_stress-VascularBundle interactions showed evidence of preferential communication, indicating that these communication patterns were supported by biologically enriched celltype pairing rather than by cell number effects alone (Fig. 4i).

Taken together, these results show that terroir-associated variation in grapevine berry tissues is reflected not only in shifts in celltype abundance, but also in coordinated relationships among specific cell populations. By integrating SEED deconvolution, BLOOM validation, correlation analysis, and cell-cell communication inference, we identify Epidermal-Hypodermal and Epidermal_stress-VascularBundle as key celltype pairs associated with regional environmental variation, thereby providing a cellular perspective for understanding terroir-associated phenotypic differences in grapevine berries.

## Discussion

Our study establishes SEED as a plant-oriented deconvolution framework that integrates reference-template construction with adaptive downstream deconvolution. In contrast to many existing approaches that rely on a fixed reference and a single deconvolution strategy, SEED directly addresses a central challenge in plant deconvolution: the strong dependence of inference accuracy on reference design, dataset context, and method-template matching. This is particularly relevant in plants, where single-cell references remain limited across species, tissues, developmental stages, and environmental conditions, and where experimental workflows can introduce additional bias through protoplast preparation or nucleus isolation.

A key finding of this study is that SEED showed its clearest advantage under the most stringent low-replication benchmark condition tested. This is meaningful in plant research, where sample numbers are often constrained by field logistics, material availability, and experimental cost. The n = 100 benchmark provided an additional comparison under a larger simulated sample size. Under this condition, the performance gap among leading methods was reduced, and method rankings varied across noise levels and evaluation metrics. This pattern suggests that, when pseudo-bulk sampling is sufficiently large, deconvolution accuracy becomes less dependent on a single dominant strategy and more influenced by the specific noise structure and evaluation criterion. In contrast, SEED showed its clearest advantage under the low-replication setting, supporting its practical value for plant transcriptome studies in which biological replication is limited.

The grapevine application further illustrates that plant deconvolution can be used not only to estimate celltype abundance, but also to uncover environmentally associated cellular structure in plant tissues. In viticulture, terroir is widely understood as a multidimensional interaction among climate, soil, cultivar, and management that shapes berry composition and wine quality. Our results suggest that part of this effect can be resolved at the level of tissue cellular composition. Specifically, SEED identified region-associated shifts in several annotated cell populations linked to outer tissue identity, stress-responsive states, and vascular function, and these patterns were further supported by BLOOM, correlation analysis, and cell–cell communication inference. Together, these results indicate that terroir-associated transcriptomic variation may be organized through coordinated changes among specific cellular populations rather than reflected only at the bulk tissue level.

At the same time, several limitations should be acknowledged. The current grapevine single-nucleus reference was derived from a limited sampling design and was intended primarily as a practical reference for deconvolution rather than as a comprehensive atlas. Likewise, the present study focuses on establishing the analytical framework and demonstrating its utility in representative plant datasets, rather than providing an exhaustive characterization of grapevine cellular diversity or a full mechanistic dissection of region-associated phenotypes. We are currently expanding the single-nucleus sampling across additional biological replicates and conditions, which will provide a stronger foundation for refining the reference and improving the interpretation of cell-state variation.

Future work will build on this framework in two main directions. First, broader and more diverse reference resources will be needed to further test the performance and transferability of SEED across plant species, tissues, and environmental contexts. Second, the terroir-associated cell populations identified here provide an entry point for deeper mechanistic investigation, including how specific cell states, intercellular communication patterns, and environmental cues jointly contribute to berry phenotype formation. Thus, rather than representing an endpoint, the present study provides a methodological and biological starting point for linking plant bulk transcriptomes to cellular heterogeneity and, ultimately, to mechanism.

## Methods

### 1. SEED framework

#### 1.1 Overview of the SEED Calculation Process

SEED was developed by adapting our previously established DECEPTICONx framework to plant transcriptomic applications. Similar to DECEPTICONx, SEED is an adaptive workflow that evaluates multiple combinations of reference-template construction and deconvolution strategies for a given dataset. SEED uses two complementary strategies—Monocle3 and BayesPrism—to construct candidate Reference Templates for downstream deconvolution.

In this study, a Reference Template is defined as a celltype-resolved reference expression matrix, in which rows correspond to genes and columns correspond to annotated cell types. These templates provide the prior expression patterns required for bulk RNA-seq deconvolution. Given single-cell RNA-seq data and bulk RNA-seq data as input, SEED first constructs multiple candidate Reference Templates from the single-cell reference, then pairs each template with a panel of deconvolution methods, and finally integrates the resulting candidate predictions to obtain robust celltype proportion estimates.

### 1.2 Generation of Candidate Predictions

Let

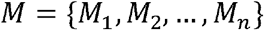

be a set of *n* deconvolution methods, and let

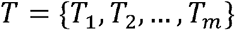

be a set of *m* candidate Reference Templates.

For a given bulk RNA-seq dataset *B*, each method–template pair (*M*_*j*_ ∣ *T*_*k*_) produces a vector of estimated proportions for all cell types. For a specific cell type *c*, the proportion estimated by pair (*M*_*j*_ ∣ *T*_*k*_) is denoted as

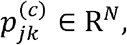

where *N* is the number of samples in *B*. Collectively, SEED obtains *n* × *m* candidate prediction vectors for each cell type. In this study, the candidate deconvolution method set consisted of CIBERSORT, quanTIseq, EPIC, DeconRNASeq, MuSiC, SCDC, and MCP-counter.

#### 1.3 Construction of candidate Reference Templates

SEED constructs candidate Reference Templates from single-cell RNA-seq data using two complementary strategies.

Monocle3: Monocle3 was used to generate celltype expression templates by grouping cells according to annotated cell identities and calculating aggregated expression profiles. In SEED, Monocle3-based preprocessing and clustering were followed by celltype-level expression summarization under different normalization schemes, thereby generating candidate reference matrices that capture aggregated expression patterns for annotated plant cell populations.

BayesPrism: BayesPrism was used to generate an alternative candidate Reference Template in a probabilistic framework. Filtered single-cell count matrices were matched to the bulk RNA-seq expression space by retaining overlapping genes between single-cell and bulk datasets, and BayesPrism was then used to construct celltype-specific reference profiles from the aligned single-cell data.

Together, these two strategies were intended to provide complementary template representations and to reduce dependence on any single reference construction method.

#### 1.4 Selection and integration of candidate predictions

Following the DECEPTICONx workflow, SEED uses the correlation structure among candidate predictions to guide result integration. For a given cell type, pairwise Pearson correlation coefficients are computed among candidate prediction vectors generated by different template– method combinations. Under the DECEPTICON framework, when the residual errors of two candidate strategies are weakly correlated, stronger agreement between their predicted proportion vectors is expected to reflect stronger agreement with the unknown true cell proportion.

If multiple candidate predictions were retained for a specific cell type, SEED assigned each selected candidate ***s*** ∈ ***S***_***c***_ an initial weight

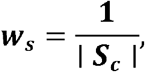

where ***S***_***c***_ is the set of selected candidates for cell type ***c***.

If a candidate appeared multiple times among top-supported combinations, its weight was increased proportionally. Let ***r***_*s*_ denote the number of times candidate ***s*** was selected. The updated weight was then defined as

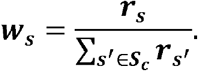

The integrated proportion estimate for cell type ***c*** across all samples was calculated as the weighted average of the selected candidate predictions:

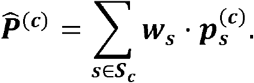

#### 1.5 Final Proportion Estimate

For each cell type, the final SEED estimate was obtained by integrating the retained candidate predictions using the weighting scheme defined above. Specifically, the final proportion estimate across all samples was calculated as the weighted sum of the selected candidate predictions:

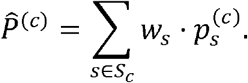

This integrated estimate was used as the final celltype proportion output for downstream analyses.

#### 1.6 Normalization and zero-value handling

For deconvolution methods whose outputs were not inherently normalized to sum to one, SEED applied sample-wise normalization:

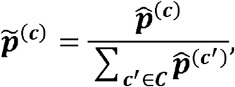

where ***c*** denotes the set of all cell types considered.

If two or more selected candidates yielded exactly zero predictions for a given cell type and their pairwise correlation was artificially inflated because of the zero values, that cell type was flagged as undetected and excluded from downstream correlation-based analyses to avoid spurious agreement.

### 2 Plant materials

Grapevine berry samples were collected during the 2023 growing season from two production regions in China: Yantai, Shandong Province, and Yinchuan, Ningxia Hui Autonomous Region. The sampled cultivar was *Vitis vinifera* cv. Cabernet Sauvignon. Berries were collected at commercial maturity.

For bulk RNA-seq sequencing, five healthy grapevines with comparable growth status and fruiting performance were selected in each production region and used as five independent biological replicates. For each biological replicate, one representative cluster was harvested together with the attached short fruiting shoot. To maintain cluster viability and minimize transcriptional changes caused by post-harvest stress, the cut end of each shoot was immediately wrapped with moist cotton and gauze. Samples were transported to the laboratory under low-temperature conditions using portable ice boxes, typically within 4–8 h after collection.

Upon arrival, samples were processed immediately. For each biological replicate, three healthy, well-colored berries were selected from the upper, middle, and lower positions of the corresponding cluster. The three berries were pooled, and seeds were carefully removed on a pre-cooled working surface. The remaining skin tissues, including skin and flesh, were immediately frozen in liquid nitrogen and stored independently at −80°C until RNA extraction.

For single-nucleus RNA sequencing, one biological sample was collected from the Yantai production region during the same sampling period as the bulk RNA-seq. One healthy grapevine with representative growth and fruiting status was randomly selected from the same vineyard block, and one representative cluster was harvested together with the attached short fruiting shoot. The sample was transported to the laboratory under the same low-temperature conditions described above. After arrival, one healthy and uniformly colored berry representing the average maturity level of the sample was selected. Seeds were removed on a pre-cooled surface, and the fresh berry skin and flesh tissues were immediately used for nuclei isolation.

### 3 Single-nucleus RNA sequencing

#### 3.1 Nuclei isolation and quality control

The fresh seedless berry sample of *Vitis vinifera* cv. Cabernet Sauvignon from Yantai was used for single-nucleus RNA sequencing. High-quality nuclei suspensions were prepared under low-temperature conditions. Briefly, fresh tissue was gently homogenized in an optimized nuclei lysis buffer and sequentially filtered through 100 μm and 40 μm cell strainers. Nuclei were enriched by centrifugation and washed with nuclei washing buffer. The purified nuclei were finally resuspended in phosphate-buffered saline containing 1% bovine serum albumin and RNase inhibitor.

Nuclei quality was assessed by DAPI staining and fluorescence microscopy to evaluate nuclei integrity, morphology, and purity. Samples with no obvious cellular debris or cytoplasmic contamination were retained. Trypan blue staining was used to confirm that intact cell contamination was below 5%. Nuclei were counted using a hemocytometer, and the final nuclei suspension was adjusted to approximately 1,000–1,200 nuclei/μL for library construction.

#### 3.2 snRNA-seq library construction and sequencing

Single-nucleus RNA-seq libraries were prepared using the 10x Genomics Chromium Next GEM Single Cell 3′ Reagent Kits v3.1 according to the manufacturer’s protocol. Qualified nuclei suspensions were loaded onto the Chromium Controller with a target recovery of approximately 10,000 nuclei. Gel bead-in-emulsion generation, reverse transcription with cell barcodes and unique molecular identifiers, cDNA amplification and purification, enzymatic fragmentation, end repair, A-tailing, adapter ligation, and sample index PCR amplification were performed following the standard 10x Genomics workflow.

The libraries were quantified using a Qubit 4.0 fluorometer, and fragment size distribution was assessed using an Agilent Bioanalyzer 2100 High Sensitivity DNA kit. Libraries with a major fragment size distribution of approximately 400–600 bp were used for sequencing.

Qualified libraries were sequenced on an Illumina NovaSeq 6000 platform using a paired-end 150 bp sequencing strategy. A total of 357,143,206 raw reads were generated. After base calling and quality filtering, 107.14 Gb of high-quality clean data were obtained. The Q20 and Q30 percentages were 96.88% and 92.01%, respectively, and the average GC content was 32.57%. The read structure followed the standard configuration of the 10x Genomics Single Cell 3′ v3.1 protocol.

### 4 Bulk RNA sequencing

#### 4.1 RNA extraction and quality control

Total RNA was extracted from mature Cabernet Sauvignon berry skin and flesh samples collected from Yantai and Yinchuan, with five biological replicates per region. Samples stored at −80°C were used for RNA extraction. Total RNA was extracted using a modified CTAB-LiCl precipitation method, followed by DNase I treatment to remove genomic DNA contamination.

RNA concentration and purity were evaluated using a NanoDrop 2000 spectrophotometer based on OD260/280 and OD260/230 ratios. RNA integrity was assessed using an Agilent Bioanalyzer 2100. Only RNA samples with an RNA integrity number of at least 7.0 were used for library construction.

#### 4.2 Bulk RNA-seq library construction and sequencing

Messenger RNA was enriched using the NEBNext Poly(A) mRNA Magnetic Isolation Module. Strand-specific RNA-seq libraries were constructed using the NEBNext Ultra II Directional RNA Library Prep Kit for Illumina according to the manufacturer’s instructions. The library construction procedure included mRNA fragmentation, cDNA synthesis with dUTP incorporation, adapter ligation, PCR amplification for 12–15 cycles, and size selection to obtain fragments of approximately 200–400 bp.

Libraries were sequenced on an Illumina NovaSeq 6000 platform using a paired-end 150 bp strategy. The target sequencing depth was at least 6 Gb of clean data per sample, with a Q30 percentage of at least 85%.

### 5 snRNA-seq data processing and celltype annotation

Raw snRNA-seq data were processed using Cell Ranger v6.1.2. The workflow included read alignment, cell barcode identification, UMI counting, and gene expression quantification. Reads were aligned to the *Vitis vinifera* cv. Cabernet Sauvignon cl.08 v1.1 (VvCabSauv08.v.1.1) reference genome (https://www.grapegenomics.com/).

Downstream analysis was performed using the R package Seurat v4.1.1. A Seurat object was created from the gene expression matrix. Genes expressed in fewer than three nuclei and nuclei with fewer than 200 detected features were removed. High-quality nuclei were retained based on the number of detected genes, total UMI counts, and the proportion of reads mapped to a customized grape mitochondrial gene set. Nuclei with mitochondrial gene expression proportions greater than 5% were excluded.

The filtered expression matrix was normalized using the LogNormalize method with a scale factor of 10,000. The top 2,000 highly variable genes were identified using the “vst” method. The data were then centered and scaled using the ScaleData function. Principal component analysis was performed, and the first 20 principal components were retained for downstream analysis. A K-nearest-neighbor graph was constructed using these principal components, and nuclei were clustered using the Louvain algorithm with a resolution of 0.3. In total, eight clusters were identified. Uniform Manifold Approximation and Projection was used for visualization based on the first 20 principal components.

Cluster marker genes were identified using the FindMarkers function with the Wilcoxon rank-sum test. The parameters were set as min.pct = 0.1, logfc.threshold > 0.25, and only.pos = TRUE. Marker genes were identified based on VvCabSauv08.v.1.1 gene IDs.

To assign biological identities to the clusters, marker genes were functionally annotated using cross-genome gene ID mapping. Coding sequences from the VvCabSauv08.v.1.1 genome were aligned to the NCBI *Vitis vinifera* cv. Pinot Noir PN40024 reference genome, which serves as the basis for the KEGG vvi database, to obtain corresponding KEGG IDs. In addition, mappings between VvCabSauv08.v.1.1 gene IDs and the GSVIV nomenclature system used in the VitisNet^49^ database were established.

Final cell-cluster annotations were determined manually by integrating multiple lines of evidence, including KEGG pathway enrichment of mapped marker genes, expression patterns of pathway genes from the VitisNet database, particularly genes related to flavonoid and anthocyanin biosynthesis, and previously reported celltype-specific marker genes from grape and related plant species. Based on this evidence, the eight clusters were annotated into biologically interpretable cell populations, including epidermal cells, parenchyma cells, and a cluster enriched for anthocyanin biosynthesis-related genes.

### 6 Bulk RNA-seq data processing

Raw bulk RNA-seq reads were first assessed using FastQC. Adapter sequences and low-quality reads were removed using Trimmomatic to obtain high-quality clean reads.

Clean reads were aligned to the VvCabSauv08.v.1.1 reference genome using HISAT2. Gene-level read counts were quantified using featureCounts from the Subread package v2.0.1 based on the corresponding genome annotation file. Strand-specific information was considered during read counting. The resulting raw count matrix was used for downstream transcriptomic analyses and SEED-based deconvolution.

### 7 Generation of simulated datasets

Simulated pseudo-bulk RNA-seq profiles were generated from the annotated grapevine single-nucleus reference dataset. Celltype-level expression signatures were first obtained by averaging raw counts across cells from each annotated cell population. For each of 100 simulated samples, random celltype fractions were generated and normalized to sum to one. Pseudo-bulk expression profiles were then constructed as weighted sums of the celltype-level expression signatures using these simulated fractions.

To assess robustness to expression noise, Gaussian perturbations were added to the simulated pseudo-bulk profiles at relative noise levels of 0, 0.1, 0.5 and 0.9, with larger values corresponding to stronger noise. Negative values after perturbation were set to zero. These datasets provided pseudo-bulk mixtures with known celltype compositions for systematic evaluation of deconvolution performance.

### 8 Correlation analysis

After normalization of the positive cell subset data derived from SEED analysis, Pearson correlation analysis was performed to evaluate the association strength between celltypes. Statistical significance was assessed by calculating P-values, with false discovery rate controlled using the Benjamini-Hochberg method for multiple comparison correction.

#### 9 Cell-cell communication analysis

Intercellular communication networks were analyzed using the PlantPhoneDB R package. Briefly, quality-controlled and normalized snRNA-seq data were imported into PlantPhoneDB, and a PlantPhoneDB object was constructed by integrating cell-cluster annotations. Ligand-receptor pairs were filtered based on the built-in PlantPhoneDB ligand-receptor interaction database, retaining only those expressed in at least 10% of cells in either the sender or receiver cell population to ensure reliability. The ligand-receptor pair database for grapevine was generated based on homologous alignment with *Arabidopsis*. To identify cell-type preferences, we computed the observed-to-expected ratio (Ro/e) for all cell-type pairs using the chi-square test. Significant ligand-receptor pairs between cell types were then identified using the average scoring approach.

## Author Contributions

Xinhao Hu, Yu Tang and Fulan Deng contributed equally to this work. Yu Tang and Guwei Tang wrote the manuscript. Xinhao Hu, Yu Tang, Fulan Deng performed data analysis. Zhiyu Chen contributed to the preliminary algorithmic research. Xinhao Hu collected the grape datasets. Xu Yan and Zengqiang Xia provided guidance on the data analysis. Henry H. Y. Tong, Xin Zou, and Jie Hao supervised the study and served as corresponding authors. All authors reviewed and approved the final manuscript. Henry H. Y. Tong secured funding for the project. Xin Zou led the conceptualization and oversaw the writing and editing process. Jie Hao directed the funding acquisition and provided overall supervision.

## Data availability

The raw sequencing data generated in this study, including both the grapevine berry single-nucleus RNA-seq data and bulk RNA-seq data, have been deposited in the Genome Sequence Archive (GSA) at the National Genomics Data Center, China National Center for Bioinformation / Beijing Institute of Genomics, Chinese Academy of Sciences. The dataset is publicly accessible under accession number GSA: CRA044013 at https://ngdc.cncb.ac.cn/gsa.

